# Creation of photocyclic vertebrate rhodopsin by single amino acid substitution

**DOI:** 10.1101/2021.12.06.471433

**Authors:** Kazumi Sakai, Yoshinori Shichida, Yasushi Imamoto, Takahiro Yamashita

**Author notes:** Corresponding Author: Takahiro Yamashita, Department of Biophysics, Graduate School of Science, Kyoto University, Kyoto 606-8502, Japan.

## Abstract

Opsins are universal photoreceptive proteins in animals and can be classified into three types based on their photoreaction properties. Upon light irradiation, vertebrate rhodopsin forms a metastable active state, which cannot revert back to the original dark state via either photoreaction or thermal reaction. By contrast, after photoreception, most opsins form a stable active state which can photo-convert back to the dark state. Moreover, we recently found a novel type of opsins whose activity is regulated by photocycling. However, the molecular mechanism underlying this diversification of opsins remains unknown. In this study, the molecular property of vertebrate rhodopsin successfully converted to the photocyclic and photoreversible properties by a single mutation at position 188. This revealed that the residue at position 188 contributes to the diversification of photoreaction properties of opsins by the regulation of the recovery from the active state to the original dark state.

## Introduction

Opsins are characterized as photosensitive G protein-coupled receptors and are universally found in diploblastic and triploblastic animals. All opsins share common structural elements including seven transmembrane domains and bind a light-absorbing chromophore, retinal, via a Schiff base linkage to Lys296 (based on the bovine rhodopsin numbering system) of opsin. Opsins function for both visual and non-visual photoreceptions and are classified into several groups based on their amino acid sequences ^1, 2^. Bovine rhodopsin is the most well-studied opsin ^3^ and it functions as a visual photoreceptive protein in the retina and binds 11-*cis* retinal in the dark. Photoisomerization of retinal to the all-trans form produces the meta II intermediate of rhodopsin, which couples with G protein. Meta II is a metastable active state and spontaneously converts to meta III ^4^. In addition, light irradiation of meta II induces the formation of meta III rather than the original dark state ^5, 6^. That is, the active state meta II very inefficiently converts back to the original dark state by photoreaction or thermal reaction. These observations show that vertebrate rhodopsin is specialized for photoactivation, and is thus characterized as a mono-stable opsin. By contrast, mollusk and arthropod rhodopsins form a stable active state, the acid-meta state, by photoisomerization of 11-*cis* to all-*trans* retinal, and the active state can photo-convert back to the original dark state, which contains 11-*cis* retinal ^2, 3^. That is, these opsins have two stable states, the dark and active states, which are reversible with each other by light irradiations and thus are known as bistable opsins. Recent accumulation of knowledge about the molecular properties of opsins revealed that many members of various opsin groups are bistable opsins (Fig. S1), which suggests that vertebrate rhodopsin evolved as a mono-stable opsin from an ancestral bistable opsin ^1^.

Recently, we identified a novel type of opsin, Opn5L1, as a photocycle opsin ^7^. Opn5L1 binds all-*trans* retinal, not 11-*cis* retinal, to form the active state in the dark. Light irradiation suppresses the G protein activation ability of Opn5L1 by the photoisomerization of the retinal to 11-cis form. Subsequent formation of a covalent adduct between the retinal and Cys188 of the opsin induces the conversion of the C11=C12 double bond to a single bond in the retinal. Afterward, the thermal rotation of the C11-C12 single bond in the retinal results in the dissociation of the Cys188-retinal adduct and the regeneration of the original dark state. The combination of photoisomerization and thermal isomerization of retinal regulates the G protein activation ability by Opn5L1, making this the first animal opsin whose ability is controlled by its photocyclic reaction.

The comparison of the amino acid sequences among opsins shows that the cysteine residue at position 188 is well-conserved in Opn5L1 group but rarely found in other opsin groups ^7, 8^, which supports the importance of Cys188 for the unique photocyclic reaction in Opn5L1. On the other hand, vertebrate rhodopsin and cone pigments, which are characterized as mono-stable opsins, share glycine residue at this position (Fig. S1). In this study, we analyzed whether or not the mutation at position 188 could change the molecular property of bovine rhodopsin to acquire the photocyclic property. Our detailed analysis revealed that G188C mutant photo-converted to the active state, meta II, which thermally recovered to the original dark state. In addition, light irradiation on meta II of G188C mutant induced the reversion to the original dark state. Therefore, G188C mutant of bovine rhodopsin exhibits the photocyclic and photoreversible property and the residue at position 188 regulates the recovery from the active state to the original dark state in vertebrate rhodopsin.

## Results and discussion

### Acquisition of photocyclic property of bovine rhodopsin G188C mutant

In the previous report, we revealed that Opn5L1 shares the cysteine residue at position 188, which underlies the photocyclic reaction of the opsin ^7^. Thus, to analyze whether or not G188C mutant of bovine rhodopsin acquires the photocyclic property, we purified G188C mutant after reconstitution with 11-*cis* retinal. However, we found that G188C mutant has much lower thermal stability than wild-type. That is, G188C mutant gradually decayed during the incubation in the dark at 37 °C (Fig. 1B), whereas wild-type was quite stable under the same condition (Fig. 1A). Thus, we improved the thermal stability of G188C mutant to analyze the detailed molecular properties of the mutant. According to previous reports ^9, 10^, we introduced two cysteine residues (N2C/D282C) into the mutant and measured the thermal decay rate during incubation in the dark at 37 °C. The time-dependent spectral changes showed that this G188C/N2C/D282C mutant decayed much more slowly than G188C mutant (Fig. 1C). Thus, we compared the spectral changes among wild-type, N2C/D282C and G188C/N2C/D282C mutant at 20 °C. Wild-type shifted the spectrum into the UV region after yellow light irradiation, which is indicative of the formation of meta II intermediate containing all-*trans*-15-*anti* retinal (Fig. S2A). Subsequently, the absorbance at around 470nm increased, which shows the transition from meta II to meta III intermediate containing all-*trans*-15-*syn* retinal ^11^. These spectral changes were also observed in N2C/D282C (Fig. 1D) as mentioned in the previous report ^9^. By contrast, G188C/N2C/D282C mutant had the absorption maximum (λmax) at 487nm and also shifted the spectrum into the UV region to form meta II by yellow light irradiation. During the subsequent incubation in the dark, the decrease of the absorbance in the UV region and the concomitant increase of the absorbance at around 485nm were observed (Fig. 1E). The analysis of the retinal configurations showed that light irradiation triggered the isomerization of the retinal to all-trans form, which converted back to the 11-cis form during the subsequent incubation in the dark (Fig. 1G). This inter-conversion of the retinal isomers can explain the spectral change of G188C/N2C/D282C mutant after light irradiation. We could observe thermal recovery of the original dark state after light irradiation also at 37 °C (Fig. 1F). In addition, G188C mutant showed thermal recovery of the absorption spectrum of the original dark state after light irradiation at 20 °C (Fig. S2B). The thermal recovery of the original dark state in G188C mutant was confirmed by the increase of the amount of 11-*cis* retinal during the incubation after light irradiation (Fig. S2C). However, we observed a smaller amount of the thermal recovery in G188C mutant (Fig. S2B) than that in G188C/N2C/D282C mutant (Fig. 1E). This was probably because meta II of G188C mutant is thermally unstable and is decomposed into all-*trans* retinal and apo-protein in addition to the reversion to the original dark state during the incubation after light irradiation, which was confirmed by the acid denaturation experiment (Fig. S2D). Altogether, these results showed that G188C mutation led to the acquisition of the ability to thermally recover the original dark state from the photoactivated state.

**Figure 1.**
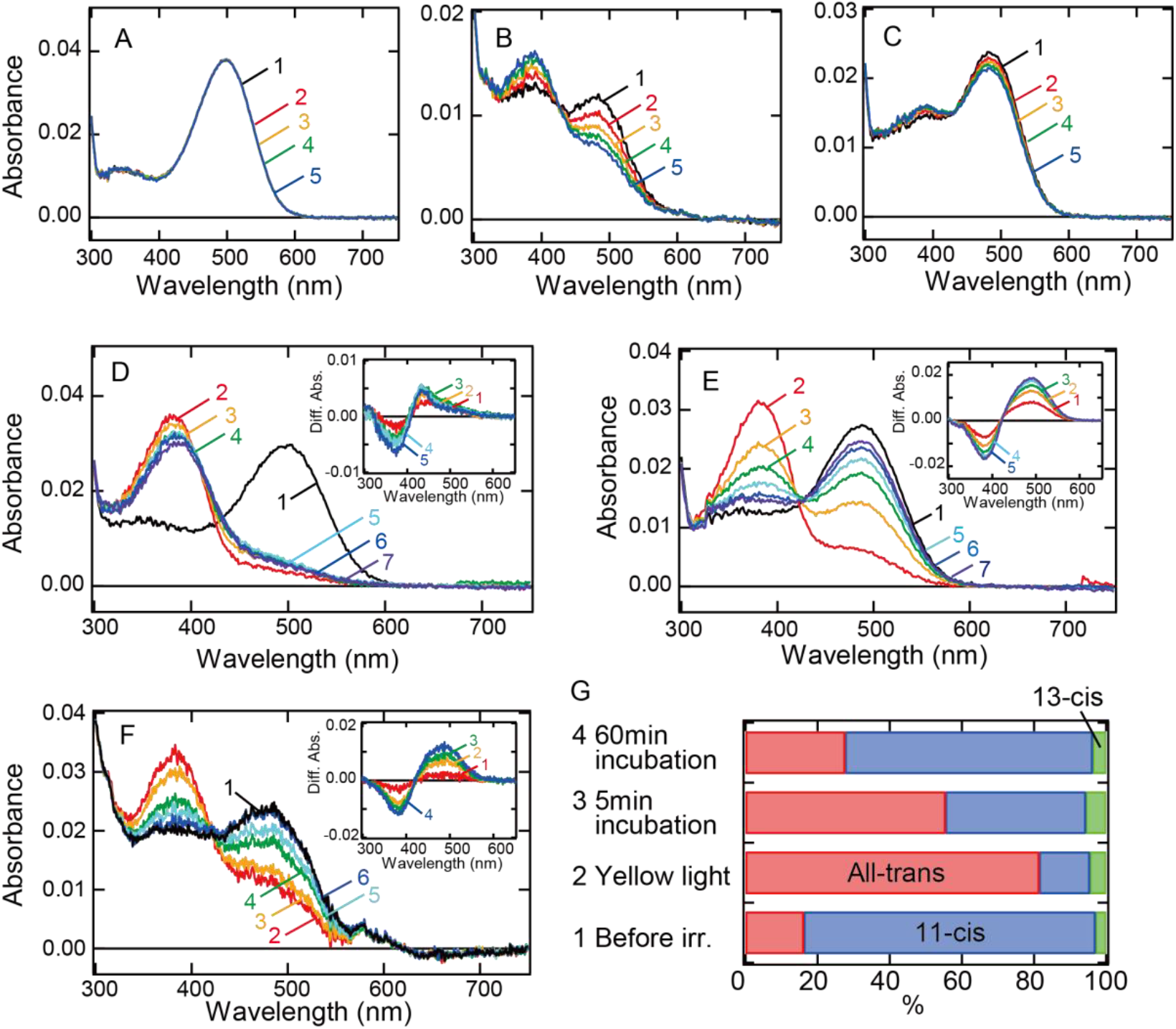
Thermal recovery of bovine rhodopsin G188C mutant after yellow light irradiation. (A-C) Thermal stability of wild-type (A) and G188C (B) and G188C/N2C/D282C (C) mutants of bovine rhodopsin purified after the incubation with 11-*cis* retinal. Absorption spectra were recorded after 0, 5, 10, 15 and 20 min incubation (curves 1-5, respectively) in the dark at 37 °C. (D, E) Absorption spectra of N2C/D282C (D) and G188C/N2C/D282C (E) mutants of bovine rhodopsin purified after the incubation with 11-*cis* retinal. Spectra were recorded in the dark (curve 1) and 0, 5, 15, 30, 60 and 120 min after yellow light irradiation (curves 2-7, respectively) at 20 °C. (Inset) Difference spectra obtained by subtracting the spectrum just after irradiation (curve 2 in Figs. 1D and 1E) from those measured after irradiation (curves 3-7 in Figs. 1D and 1E) (curves 1-5, respectively). (F) Absorption spectra of G188C/N2C/D282C mutant measured at 37 °C. Spectra were recorded in the dark (curve 1) and 0.1, 10, 50, 100 and 1000 sec after yellow flash light irradiation (curves 2-6, respectively). (Inset) Difference spectra obtained by subtracting the spectrum just after irradiation (curve 2 in Fig. 1F) from those measured after irradiation (curves 3-6 in Fig. 1F) (curves 1-4, respectively). (G) Isomeric compositions of retinal of G188C/N2C/D282C mutant. The retinal configurations were analyzed by HPLC after extraction of the chromophore from the samples before light irradiation and 0, 5 and 60 min after yellow light irradiation at 20 °C as shown in Fig. S5A.

We also analyzed whether or not other G188 mutants acquire the photocyclic property. We performed systematic mutational analysis at this position of bovine rhodopsin. A previous study showed that G188E and G188R mutants of human rhodopsin cannot form the photo-pigments after reconstitution with 11-*cis* retinal ^12^. Thus, we introduced 16 other mutations at position 188 of bovine rhodopsin and prepared the mutant proteins purified after reconstitution with 11-*cis* retinal. We successfully detected the photo-pigments from 8 of these mutants (curve 1 in Fig. S3). λmax of the mutants was blue-shifted from that of wild-type (500nm) with one exception, G188D (509nm) (Table S1). Yellow light irradiation of these mutants shifted the spectra into the UV region to form meta II (curve 2 in Fig. S3). During the subsequent incubation in the dark at 20 °C, each mutant showed characteristic spectral changes (curves 3-8 in Fig. S3). However, these spectral changes were different from a substantial increase of the absorbance at around their λmax. These results showed that the thermal recovery to the original dark state after light irradiation was not clearly detected in these mutants. Thus, we concluded that the photocyclic property was observed uniquely in G188C mutant.

### Acquisition of photoreversible property of bovine rhodopsin G188C mutant

We also analyzed whether or not meta II of G188C mutant converts back to the original dark state in a light-dependent manner. We cooled wild-type and G188C mutant to 0 °C to prevent the thermal reaction of meta II and measured their spectral changes induced by yellow light and subsequent UV light irradiations. Yellow light irradiation of wild-type resulted in the formation of meta II, and subsequent UV light irradiation shifted the spectrum into the visible region with λmax (∼470nm) blue-shifted from that of the original dark state (Fig. 2A). The previous studies revealed that this state is comparable to meta III ^5, 6^. We also constructed template absorption spectra of the dark state, meta II and mea III modeled by the Lamb and Govardovskii method ^13, 14^ (Fig. S4B) and fitted the difference spectrum (curve 2 in the inset of Fig. 2A) calculated by subtracting the spectrum after yellow light irradiation from that after UV light irradiation. Our fitting analysis showed that meta III was formed much more efficiently than the original dark state by UV light irradiation of meta II (Fig. S4C and Table S1). In addition, we analyzed the change of the retinal configuration during this process (Fig. 2C). Yellow light irradiation produced a large amount of all-*trans* retinal and completely abolished the formation of 11-*cis* retinal. Moreover, subsequent UV light irradiation produced a very limited amount of 11-*cis* retinal, which is consistent with the fitting analysis of the difference spectrum. These results confirmed that UV light irradiation of meta II induces the syn/anti isomerization of the C=N double bond of the Schiff base more efficiently than the cis/trans isomerization of the retinal. This is in contrast with the efficient cis/trans isomerization induced by the first yellow light irradiation.

**Figure 2.**
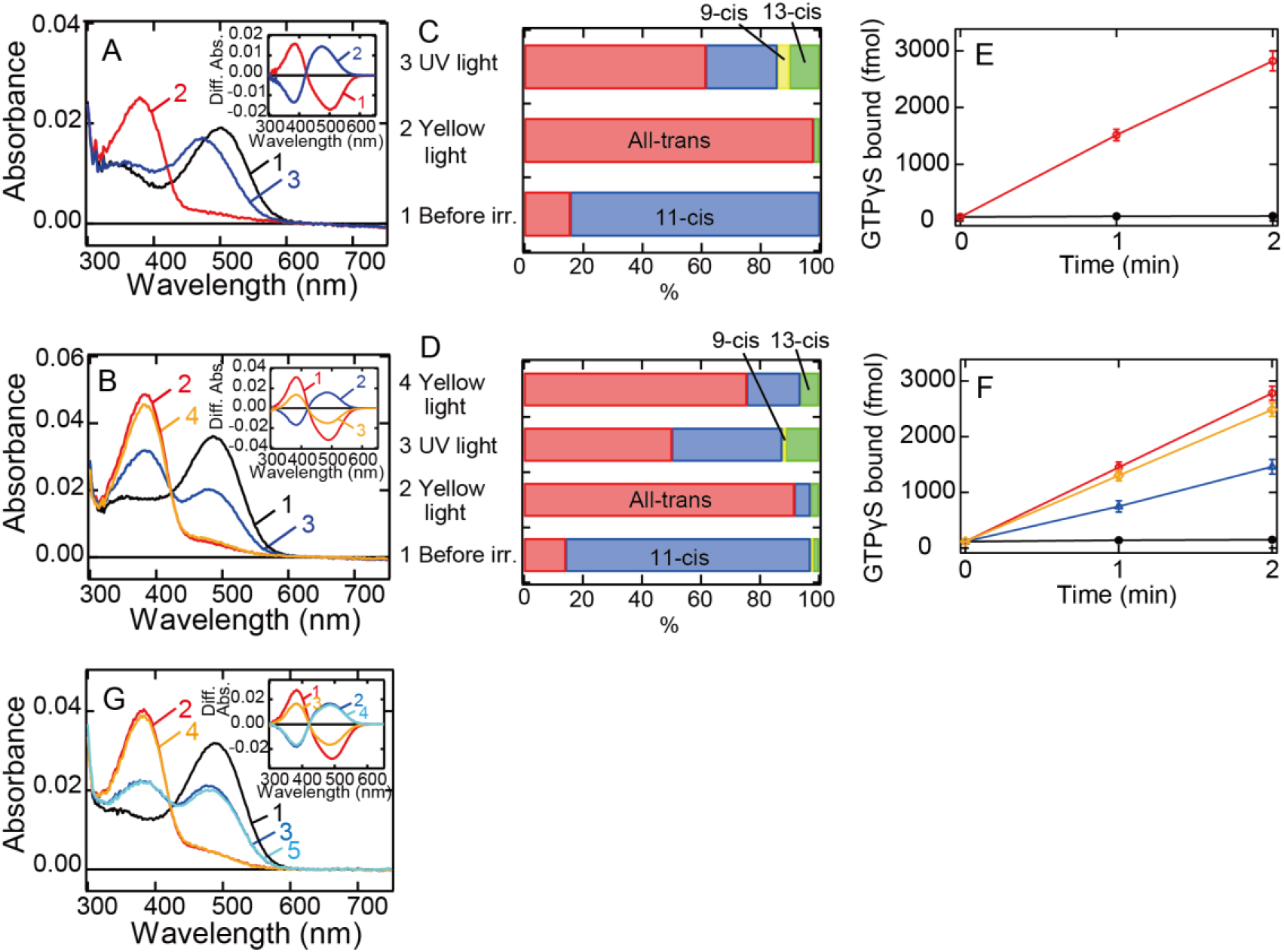
Photoreaction, retinal configuration and G protein activation of bovine rhodopsin G188C mutant. (A, B) Absorption spectra of wild-type (A) or G188C mutant (B) of bovine rhodopsin purified after the incubation with 11-*cis* retinal at 0 °C. Spectra were recorded in the dark (curve 1), after yellow light (>500 nm) irradiation (curve 2), after subsequent UV light (360 nm) irradiation (curve 3) and after yellow light re-irradiation (curve 4). (Inset) Spectral changes of wild-type (A) or G188C mutant (B) induced by yellow light irradiation (curve 1), subsequent UV light (curve 2) irradiation and yellow light re-irradiation (curve 3). Difference spectra were calculated based on the spectra shown in Figs. 2A and 2B. (C, D) Isomeric compositions of retinal of wild-type (C) and G188C mutant (D). The retinal configurations were analyzed by HPLC after extraction of the chromophore from the samples before light irradiation, after yellow light irradiation, after subsequent UV light irradiation and after yellow light re-irradiation at 0 °C as shown in Figs. S5B and S5C. (E) Gi-type of G protein activation ability by wild-type. The activation ability was measured in the dark (closed circle) and after yellow light irradiation (open circle). (F) Gi-type of G protein activation ability by G188C mutant. The activation ability was measured in the dark (closed circles), after yellow light irradiation (open circles), after subsequent UV light irradiation (open triangles) and after yellow light re-irradiation (open diamonds). Data shown in (E) and (F) were obtained at 0°C and are presented as the means ± S.E.M of three independent experiments. (G) Absorption spectrum of G188C/N2C/D282C mutant purified after the incubation with 11-*cis* retinal at 0 °C. Spectra were recorded in the dark (curve 1), after yellow light (>500 nm) irradiation (curve 2), after subsequent UV light (360 nm) irradiation (curve 3), after yellow light re-irradiation (curve 4) and after UV light re-irradiation (curve 5). (Inset) Spectral changes induced by yellow light irradiation (curve 1), subsequent UV light (curve 2) irradiation, yellow light re-irradiation (curve 3) and UV light re-irradiation (curve 4). Difference spectra were calculated based on the spectra shown in Fig. 2G.

Yellow light irradiation of G188C mutant induced the formation of meta II, and subsequent UV light irradiation shifted the spectrum into the visible region with λmax quite similar to that of the original dark state (Fig. 2B). Yellow light re-irradiation caused formation of a state whose spectrum almost overlapped that induced by the first yellow light irradiation (curve 4 in Fig. 2B). Spectral changes induced by UV light irradiation and yellow light re-irradiation are mirror images of each other (curves 2 and 3 in the inset of Fig. 2B). We fitted the UV light-dependent spectral change with the template spectra and showed that the original dark state was formed much more efficiently than meta III by UV light irradiation of meta II (Fig. S4C and Table S1). This is supported by the analysis of the retinal configurations of G188C mutant, which showed that, after the conversion from 11-*cis* to all-*trans* retinal by yellow light irradiation, subsequent UV light irradiation increased the amount of 11-*cis* retinal more efficiently than UV light irradiation of wild-type (Fig. 2D). These results suggested that meta II of G188C mutant can efficiently photo-convert back to the original dark state. Next, we measured the ability of G188C mutant to activate Gi-type of G protein, because bovine rhodopsin can activate not only transducin but also Gi/Go-types of G protein ^15, 16^. Our GTPγS binding assay showed that the light-dependent Gi activation ability of G188C was equivalent to that of wild-type (Figs. 2E and 2F). Subsequent UV light irradiation of G188C mutant suppressed the ability and yellow light re-irradiation increased the ability (Fig. 2F), which can be explained by the changes of the absorption spectra and the retinal configurations (Figs. 2B and 2D). In addition, G188C/N2C/D282C mutant could also exhibit inter-convertibility between the original dark state and meta II upon yellow light and UV light irradiations at 0 °C (Fig. 2G). These data showed that G188C mutant acquires the property of photoreversibility between the dark state and meta II. We also analyzed the photoreaction of 8 other mutants. In all of these mutants, we observed the spectral shift to the UV region by yellow light irradiation and the re-increase of the absorbance in the visible region by subsequent UV light irradiation (Figs. S4A and S4C). The spectral fitting of the difference spectra calculated before and after UV light irradiation using template spectra of the dark state, meta II and meta III (cyan curves in Fig. S4C) provided information about the component ratios of the dark state, meta II and meta III after UV light irradiation (Table S1). These results indicated that the recovery to the original dark state upon UV light irradiation occurs most efficiently in G188C mutant.

### Speeding up of the photocycle in G188C mutant

Next, we investigated whether or not the alteration of the lifetime of meta II can modulate the photocycle rate of G188C mutant. It has been reported that E122Q mutation of vertebrate rhodopsin accelerates the decay of meta II to shorten the lifetime of meta II ^17, 18^. Thus, we prepared E122Q/G188C/N2C/D282C mutant and measured its spectral change after light irradiation. Our spectral and retinal configuration analyses confirmed the thermal recovery of the original dark state in E122Q/G188C/N2C/D282C mutant after light irradiation at 0 °C (Figs. 3A and 3B). In addition, the photocycle rate of E122Q/G188C/N2C/D282C mutant at 37 °C (Fig. 3C) was about 12 times faster than that of G188C/N2C/D282C mutant (Fig. 3D). Thus, the alteration of the lifetime of meta II by single mutation successfully speeded up the photocycle of G188C mutant.

**Figure 3.**
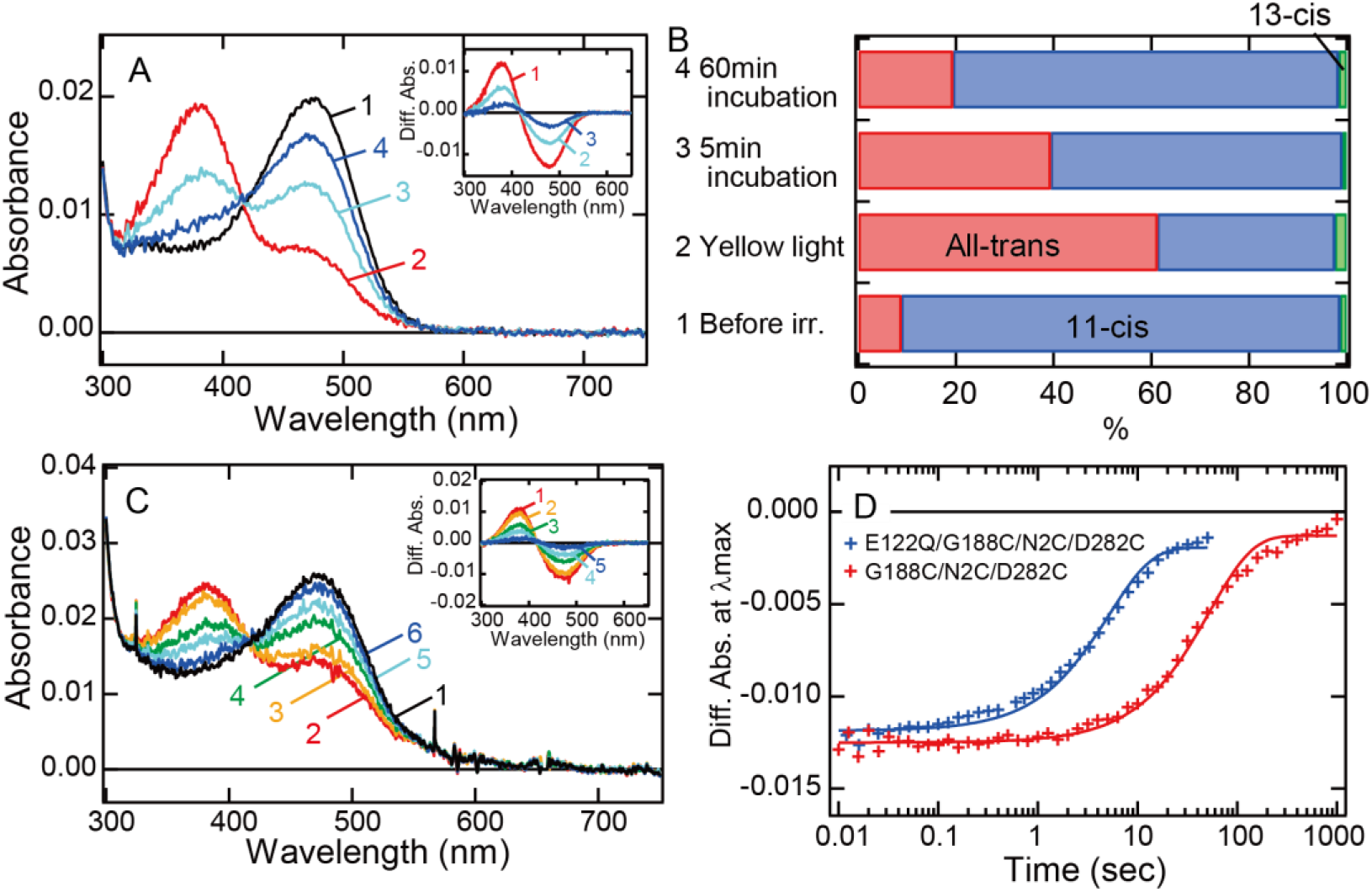
Faster recovery rate of the photocycle of bovine rhodopsin G188C mutant by introducing E122Q mutation. (A) Absorption spectra of E122Q/G188C/N2C/D282C mutant measured at 0 °C. Spectra were recorded in the dark (curve 1) and 0, 5 and 60 min after yellow light (>500 nm) irradiation (curves 2-4, respectively). (Inset) Difference spectra obtained by subtracting the spectrum before irradiation (curve 1 in Fig. 4A) from those measured after irradiation (curves 2-4 in Fig. 3A) (curves 1-3, respectively). (B) Isomeric compositions of retinal of E122Q/G188C/N2C/D282C mutant. The retinal configurations were analyzed by HPLC after extraction of the chromophore from the samples before light irradiation and 0, 5 and 60 min after yellow light irradiation at 0 °C as shown in Fig. S5D. (C) Absorption spectra of E122Q/G188C/N2C/D282C mutant measured at 37 °C. Spectra were recorded in the dark (curve 1) and 0.1, 1, 5, 10 and 50 sec after yellow flash light irradiation (curves 2-6, respectively). (Inset) Difference spectra obtained by subtracting the spectrum before irradiation (curve 1 in Fig. 3C) from those measured after irradiation (curves 2-6 in Fig. 3C) (curves 1-5, respectively). (D) Comparison of the thermal recovery process between G188C/N2C/D282C (red) and E122Q/G188C/N2C/D282C (blue). Difference absorbance at λmax obtained by subtracting the spectrum before irradiation from those measured after irradiation shown in Figs. 1F and 3C was plotted against time elapsed after irradiation. The time constants of the thermal recovery to the dark state of G188C/N2C/D282C and E122Q/G188C/N2C/D282C mutants at 37 °C were 57.4 sec and 5.1 sec, respectively.

### Modulation of G protein activation ability by the photocyclic property

We also investigated whether or not the acquisition of the photocyclic property by G188C mutation affects the G protein activation ability. As shown in Fig. 2, light-dependent Gi activation ability was equivalent between wild-type and G188C mutant at 0 °C, where substantial thermal recovery to the original dark state was not observed in G188C mutant. Thus, we measured the intracellular cAMP level in the cultured cells using cAMP biosensor (GloSensor) and compared the change of the luminescence from the biosensor triggered by bovine rhodopsin. The increase of the cAMP level induced by the addition of forskolin was attenuated by yellow light irradiation in the N2C/D282C bovine rhodopsin-transfected cells (Figs. 4A and 4B), but not in the mock-transfected cells (Fig. 4G), and it subsequently recovered slowly. By contrast, in the G188C/N2C/D282C mutant-transfected cells, we observed rapid recovery of the cAMP level after the fall of the level induced by yellow light irradiation (Figs. 4C and 4D). In addition, in the E122Q/G188C/N2C/D282C mutant-transfected cells, the cAMP level recovered more quickly from the decline induced by yellow light irradiation (Figs. 4E and 4F). These results showed that the acquisition of the photocyclic property by G188C mutation changes the G protein activation profile by promoting fast recovery to the original dark state.

**Figure 4.**
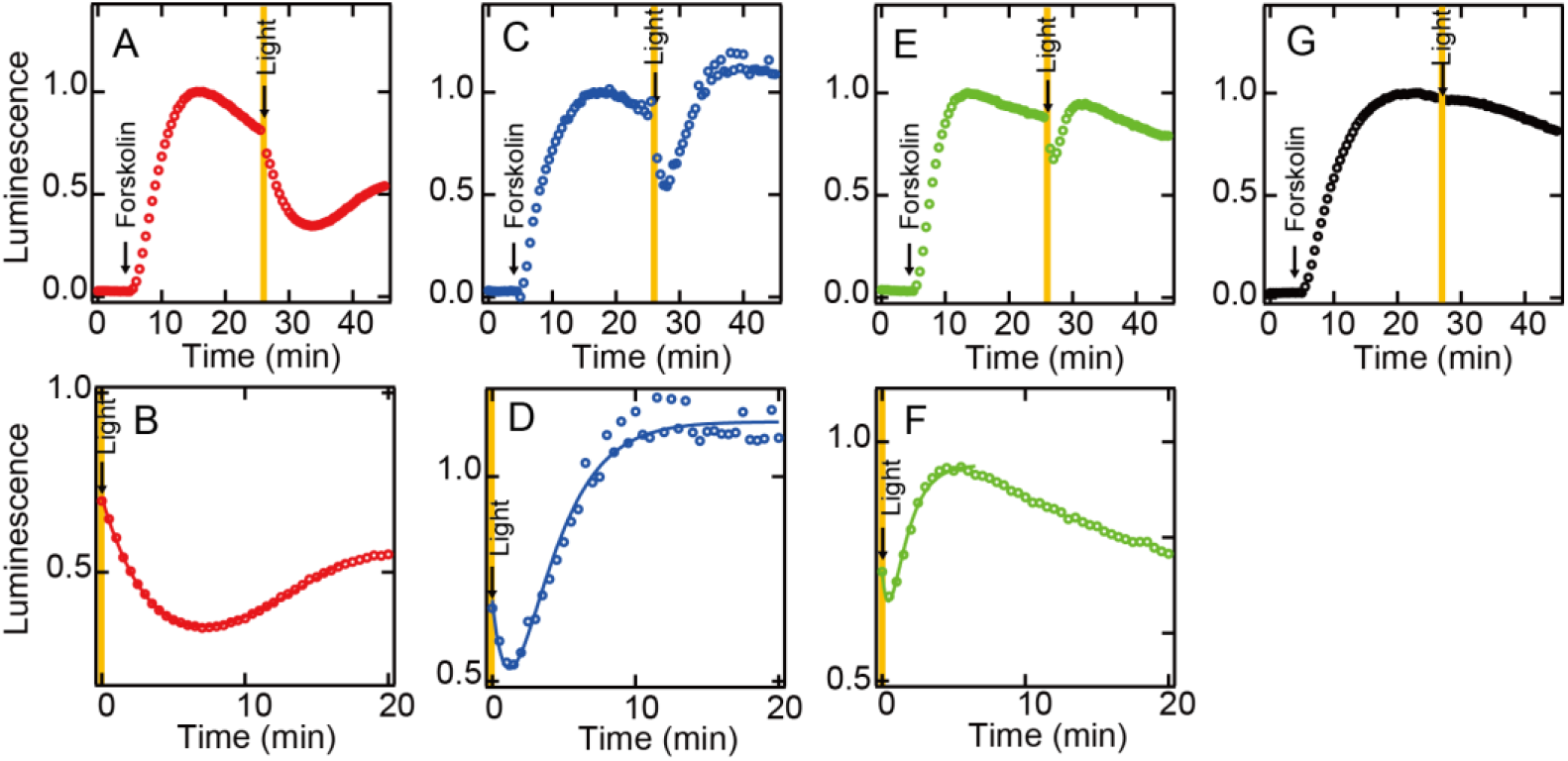
Light-mediated suppression of intracellular cAMP level by bovine rhodopsin mutants. The cAMP levels in N2C/D282C-(A, B), G188C/N2C/D282C-(C, D), E122Q/G188C/N2C/D282C-(E, F) and mock-(G) transfected HEK293T cells were measured using GloSensor cAMP assay at room temperature. The cells were incubated with 5 µM 11-*cis* retinal for 2 hr and subsequently treated with 2 µM forskolin prior to the exposure to yellow light (>500 nm). Data were normalized to the maximum point before light irradiation. Detailed profiles of the light-dependent cAMP level changes in N2C/D282C, G188C/N2C/D282C and E122Q/G188C/N2C/D282C are shown in (B), (D) and (F), respectively.

### Formation of the photo-pigments by G188C mutant upon the addition of all-*trans* retinal

We finally analyzed whether or not G188C mutant forms the photo-pigments after reconstitution with all-*trans* retinal. We purified wild-type and G188C mutant after the addition of all-*trans* retinal to the suspension of rhodopsin-expressing cell membranes. The absorption spectrum of wild-type in the dark had almost no peaks in the visible and near-UV regions (Fig. S6A). By contrast, the absorption spectrum of G188C mutant in the dark had a peak in the visible region (Fig. S6B), which was derived from predominant incorporation of 11-*cis* and 9-*cis* retinals, not all-*trans* retinal (Fig. S6C). Yellow light irradiation of this pigment resulted in the conversion of the retinal to all-trans form to shift the spectrum into the UV region, and subsequent UV light irradiation re-increased the absorbance in the visible region by the isomerization of the retinal from the all-trans to 11-cis form at 0 °C (Figs. S6B and S6C). This is quite similar to the finding for G188C mutant purified after reconstitution with 11-*cis* retinal (Figs. 2B and 2D).

We also prepared the purified apo-proteins of N2C/D282C and G188C/N2C/D282C and investigated the regeneration process of the photo-pigments upon the addition of 11-*cis* or all-*trans* retinal. The addition of 11-*cis* retinal to N2C/D282C and G188C/N2C/D282C quickly increased the absorbance at around 505 nm (Figs. S6D and S6E) and 490 nm (Figs. S6H and S6I), respectively, which showed the formation of their 11-*cis* retinal bound dark states. The addition of all-*trans* retinal to N2C/D282C resulted in a slight increase of the absorbance at around 480 nm (Figs. S6F and S6G), whereas the addition of all-*trans* retinal to G188C/N2C/D282C resulted in a substantial increase of the absorbance at around 485 nm (Figs. S6J and S6K). The regeneration ability of G188C/N2C/D282C in response to the addition of all-*trans* retinal was much higher than that of N2C/D282C (Fig. S6L). These results showed that G188C mutant can uniquely form the photo-pigments upon the addition of not only 11-*cis* retinal but also all-*trans* retinal.

### Mechanism of conversion of the molecular property by G188C mutation

In general, vertebrate rhodopsin photo-converts to a metastable active state, meta II, by the cis/trans isomerization of the retinal and subsequently undergoes a thermal transition to meta III by the syn/anti isomerization of the C=N double bond of the Schiff base ^6, 11, 19^. Thus, in response to light and heat, meta II exhibits greater conversion from all-*trans*-15-*anti* to all-*trans*-15-*syn* retinal than from all-*trans* to 11-*cis* retinal and recovers to the original dark state very inefficiently. By contrast, G188C mutant can revert to the original dark state from meta II much more efficiently than wild-type, which possibly results from the preferential isomerization from all-*trans* to 11-*cis* retinal within the chromophore binding pocket of meta II. The molecular model of G188C mutant meta II constructed based on the crystal structure of wild-type meta II ^20^ suggests that Cys188 can be located in the vicinity of Glu181 (Fig. S7). Glu181 works as a counterion to stabilize the protonation of the Schiff base in meta I, a precursor of meta II ^21, 22^. It should be noted that G188D mutant formed a substantial amount of meta I after light irradiation (Fig. S3A), which can be explained by the stabilization of the protonated Schiff base of meta I by the introduction of the aspartic acid residue. Thus, the introduction of cysteine residue at position 188 may affect the local structure and the chemical environment around the Schiff base, which prevents the syn/anti isomerization of the C=N double bond of the Schiff base and accelerates the cis/trans isomerization of the retinal.

The addition of all-*trans* retinal to G188C mutant resulted in the formation of 9-*cis* or 11-*cis* retinal-containing photo-pigments (Fig. S6). 9-*cis* or 11-*cis* retinal within G188C mutant would be formed from all-*trans* retinal on the outside or inside of the protein. If the thermal isomerization of the retinal occurs on the outside of the protein, we can expect the formation of the photo-pigments also from wild-type. However, this was observed much less efficiently in wild-type. Thus, we speculate that 9-*cis* or 11-*cis* retinal would be formed from all-*trans* retinal on the inside of G188C mutant, which is consistent with the acceleration of the thermal cis/trans isomerization of the retinal in meta II of G188C mutant.

We observed the thermal recovery from the photoactivated state in G188C mutant (Figs. 1 and S2), but not in other mutants (Fig. S3). Thus, the cysteine residue introduced at position 188 would have a special role to facilitate the thermal isomerization of the retinal. In a previous study of Opn5L1, we revealed that the light-dependent adduct formation between Cys188 and 11-*cis* retinal accelerates the isomerization to all-*trans* retinal to recover the original dark state ^7^. During this photocycle process of Opn5L1, our spectral analysis detected an increase of the absorbance at around 270 nm (Figs. S8A and S8B), which is derived from the breaking of the retinal-conjugated double bond system by the adduct formation. However, in this study, the introduction of the cysteine residue at position 188 of bovine rhodopsin induced the isomerization from all-*trans* to 11-*cis* retinal, which is a reverse reaction to the thermal isomerization of the retinal in Opn5L1. Moreover, we could not clearly observe an increase of the absorbance at around 270 nm during the thermal reaction of bovine rhodopsin G188C mutant after photoreception (Figs. S8C and S8D). Thus, the detailed molecular mechanism of the acceleration of the thermal recovery to the original dark state in G188C mutant remains unknown, but we speculate that the cysteine residue introduced at position 188 possibly transiently forms an adduct with all-*trans* retinal after photoactivation and quickly dissociates from the retinal after the isomerization to 11-*cis* retinal.

## Conclusion

In this study, we analyzed a series of mutants at position 188 of bovine rhodopsin and found that G188C mutant has a unique active state which can revert to the original dark state both by a thermal reaction and in a light-dependent manner. These results showed that the molecular property of vertebrate rhodopsin can convert to photocyclic and photoreversible properties by this single mutation. Little attention has been paid to the functional role of the residue at position 188 in opsins so far. The combination of the mutation at position 188 with other mutations in various opsins could perturb the local structure around the Schiff base, which could lead to inter-conversion of the molecular properties among opsins. Moreover, G188C mutant of bovine rhodopsin has several advantages as an optogenetic tool, because G188C mutant can be reconstituted in the presence of all-*trans* retinal and exhibits the photocyclic property, like channelrhodopsin ^23^ in addition to its high expression yield in mammalian cultured cells and high G protein activation ability. We successfully showed that the change of the lifetime of meta II by the single mutation can modulate the photocycle rate of G188C mutant. Accumulation of evidence about the mutants of vertebrate rhodopsin can help to guide the modification of the molecular properties of G188C mutant, which will provide a novel type of optogenetic tools based on vertebrate rhodopsin.

## Materials and Methods

### Preparation of bovine rhodopsin mutants

The mutant cDNAs of bovine rhodopsin (accession no. AB062417) were constructed using an In-Fusion cloning kit (Clontech). The wild-type and mutant cDNAs of bovine rhodopsin were inserted into the mammalian expression vector pUSRα ^24^ or pCAGGS ^25^. The plasmid was transfected into HEK293T cells using the calcium-phosphate method. After 2 days incubation, the transfected cells were collected by centrifugation and suspended in Buffer A (50 mM HEPES, 140 mM NaCl, pH 6.5), and 11-*cis* or all-*trans* retinal was added to the cell suspension to reconstitute the photo-pigments. They were solubilized in Buffer A containing 1% dodecyl maltoside (DDM) and adsorbed to a Rho1D4 (the anti-bovine rhodopsin monoclonal antibody) affinity column to purify the pigments. After washing the column with Buffer A containing 0.02 % DDM, the pigment was eluted by adding synthetic peptide with the epitope sequence. To purify the apo-proteins of rhodopsin, the transfected cell membranes without the addition of the retinal were solubilized in Buffer A containing 1% DDM and adsorbed to a Rho1D4 affinity column.

### Spectroscopic measurements

UV/Vis absorption spectra were recorded with a UV-visible spectrophotometer (UV-2450 and UV-2400, Shimadzu). Samples were kept at 0, 20 or 37 °C using a cell holder equipped with a circulation system that uses temperature-controlled water in order to analyze the thermal reaction of the pigments in detail. The samples were irradiated with either yellow light through a Y-52 cutoff filter (Toshiba) or UV light through a UVD-36 glass filter (AGC Techno Glass) from a 1 kW tungsten halogen lamp (Master HILUX-HR; Rikagaku).

To measure the photocycle process of G188C mutant of bovine rhodopsin, a time-resolved CCD spectrophotometer (C10000 system, Hamamatsu Photonics) was used ^26^. Spectra were taken from G188C mutant samples in the dark and at different time points after irradiation (170-μs, yellow light through a Y-52 cutoff filter from a Xenon flash lamp). Temperature of the sample was kept at 37 °C by a temperature controller (pqod, QUANTUM Northwest). Absorbance changes at λmax were plotted as a function of time and fitted with a single-exponential function to obtain the time constants for the recovery to the original dark state.

### Retinal configuration analysis

Retinal configurations within rhodopsin samples were analyzed by HPLC (LC-10ATvp; Shimadzu) equipped with a silica column (YMC-Pack SIL, particle size 3 μm, 150 × 6.0 mm, YMC) as previously described ^27^.

### G protein activation assay

The activation of Gi-type of G protein was measured by GDP/GTPγS exchange of G protein using a radionucleotide filter-binding assay ^15, 28^. Giαβγ was prepared by mixing rat Giα1 expressed in E. coli strain BL21 ^29^ with Gtβγ purified from bovine retina ^30^. All of the assay procedures were carried out at 0 °C. The assay mixture consisted of 10 nM pigment, 600 nM G protein, 50 mM HEPES (pH 7.0), 140 mM NaCl, 5 mM MgCl_2_, 1 mM DTT, 0.01% DDM, 1 μM [^35^S]GTPγS and 2 μM GDP. Bovine rhodopsin wild-type and G188C mutant purified after reconstitution with 11-*cis* retinal were mixed with G protein solution and were kept in the dark or irradiated with yellow light (>500 nm) for 1 min, with subsequent UV light for 1 min or with yellow light re-irradiation for 1 min. After irradiation, the GDP/GTPγS exchange reaction was initiated by the addition of [^35^S]GTPγS solution to the mixture of rhodopsin and G protein. After incubation for the selected time in the dark, an aliquot (20 μl) was removed from the sample into 200 μl of stop solution (20 mM Tris/Cl (pH 7.4), 100 mM NaCl, 25 mM MgCl_2_, 1 μM GTPγS and 2 μM GDP), and it was immediately filtered through a nitrocellulose membrane to trap [^35^S]GTPγS bound to G proteins. The amount of bound [^35^S]GTPγS was quantitated by assaying the membrane with a liquid scintillation counter (Tri-Carb 2910 TR; PerkinElmer).

### cAMP level measurement in cultured cells

cAMP levels in HEK293T cells were measured using the GloSensor cAMP assay (Promega) according to the manufacturer’s instructions and a previous report ^31^. HEK293T cells were seeded in 96-well plates at a density of 20,000 cells/well in low serum medium (D-MEM/F12 containing 0.25 % FBS). After 24 hours of incubation, cells were transfected with 50ng of rhodopsin plasmid and 50 ng of Glosensor 22F plasmid per well by the polyethylenimine transfection method. After overnight incubation, the medium was replaced with an equilibration medium which contained a 2% dilution of the GloSensor cAMP reagent stock solution, 10 % FBS and 5 µM retinal in a CO_2_-independent medium (Thermo Fisher Scientific). Following 2 hours equilibration at room temperature, luminescence from the cells was measured using a microplate reader (SpectraMax L, Molecular Devices). For the measurement of Gi activation by wild-type and mutant rhodopsin, the cells were first treated with 2 µM forskolin to increase the cAMP-dependent luminescence to the plateau level and subsequently stimulated for 30 sec with yellow light through a Y-52 cutoff filter from a 1 kW tungsten halogen lamp.

## Acknowledgements

We thank Dr. E. Nakajima for a critical reading of the manuscript. We also thank Prof. R. S. Molday for the generous gift of a Rho1D4-producing hybridoma and Dr. K. Sato for valuable discussion of the manuscript and technical advice about the cAMP assay.

## Author Contributions

K.S., Y.S., and T.Y. designed research; K.S. and T.Y. performed research; Y.I. contributed new reagents and analytic tools; K.S., Y.S., and T.Y. analyzed data; K.S., Y.S., and T.Y. wrote the manuscript with editing by all authors.

## Supplementary information

### Supplementary Figures

**Figure S1.**
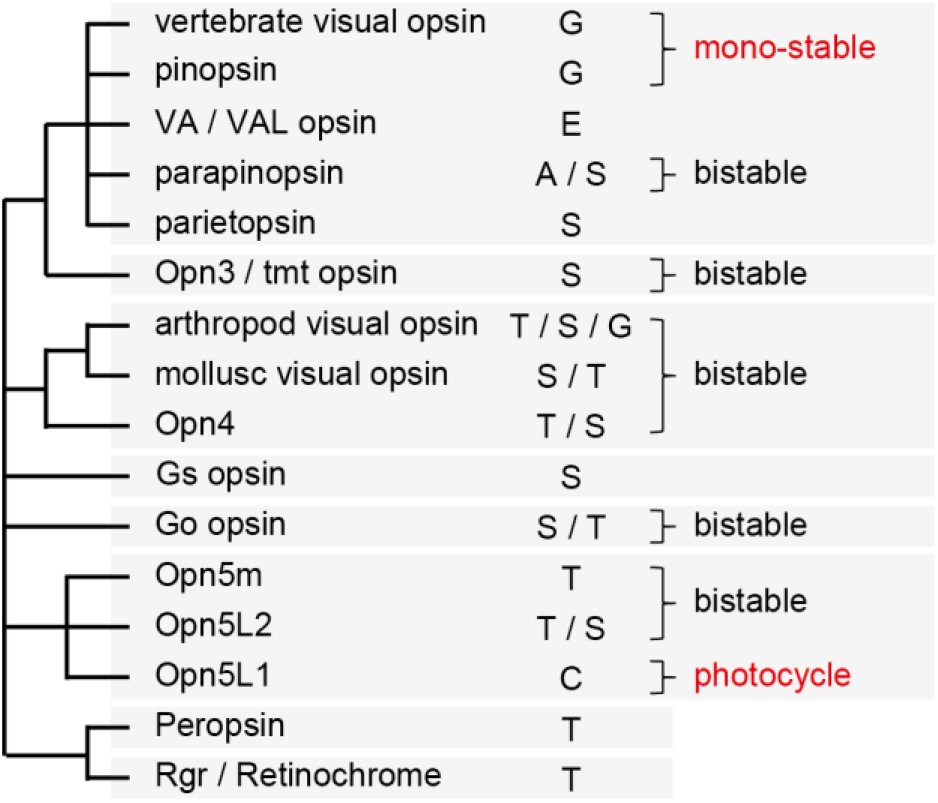
Comparison of the amino acid residue at position 188 of opsins in their phylogenetic relationship. Opn5L1 is characterized as photocycle opsin and uniquely has cysteine residue at position 188. By contrast, vertebrate visual pigments and pinopsin are characterized as mono-stable opsins and share glycine residue at position 188.

**Figure S2.**
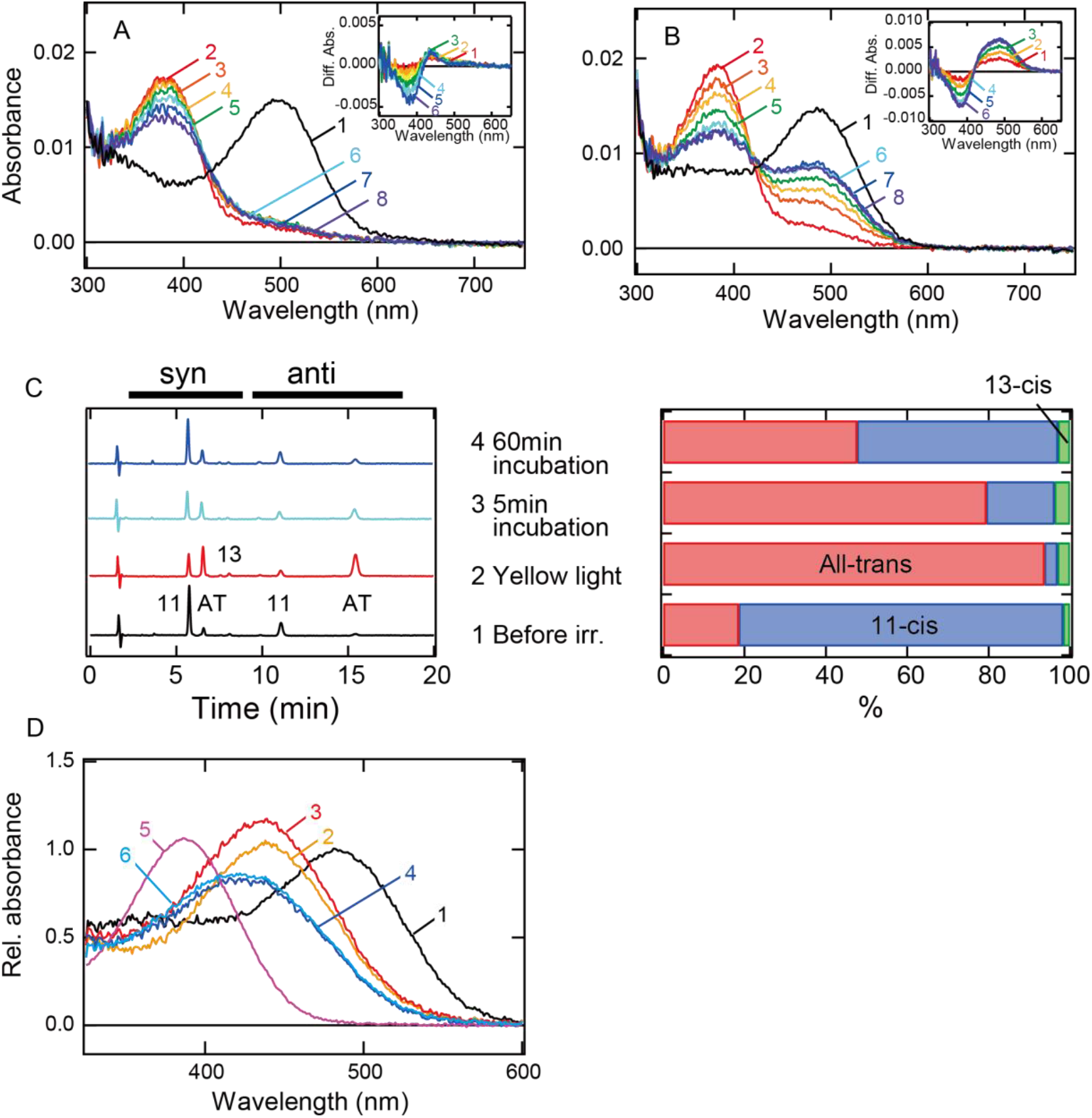
Thermal reaction of bovine rhodopsin wild-type and G188C mutant after yellow light irradiation. (A, B) Absorption spectra of wild-type (A) and G188C mutant (B) measured at 20 °C. Spectra were recorded in the dark (curve 1) and 0, 5, 10, 20, 40, 80 and 160 min after yellow light (>500 nm) irradiation (curves 2-8, respectively). (Inset) Difference spectra obtained by subtracting the spectrum just after irradiation (curve 2 in Figs. S2A and S2B) from those measured after irradiation (curves 3-8 in Figs. S2A and S2B) (curves 1-6, respectively). (C) Isomeric compositions of retinal of G188C mutant. The retinal configurations were analyzed by HPLC after extraction of the chromophore from the samples before light irradiation and 0, 5 and 60 min after yellow light irradiation at 20 °C. (D) Acid denaturation of G188C mutant samples in the dark and after yellow light (>500 nm) irradiation. Spectra of G188C mutant were recorded in the dark (curve 1) and after acid denaturing by the addition of HCl (final pH: 1.3 ± 0.2, curve 2). Spectrum recorded in the dark was normalized to be ∼1.0 at λmax. Moreover, spectra of G188C mutant were recorded after acid denaturing by the addition of HCl to meta II formed just after yellow light irradiation (curve 3) and to the original dark state converted from meta II during 60 min incubation after yellow light irradiation (curve 4). Absorption spectrum of all-*trans* retinal was also recorded in Buffer A containing 0.02 % DDM and HCl (curve 5). Curve 6 was obtained by the assumption that the acid-denatured sample shown by the curve 4 contains the acid-denatured original dark state (11-*cis* retinal bound state, curve 2) and free all-*trans* retinal (curve 5) based on the component ratio of 11-*cis* and all-*trans* retinals shown in Fig. S2C. The calculated curve 6 fitted well with the curve 4.

**Figure S3.**
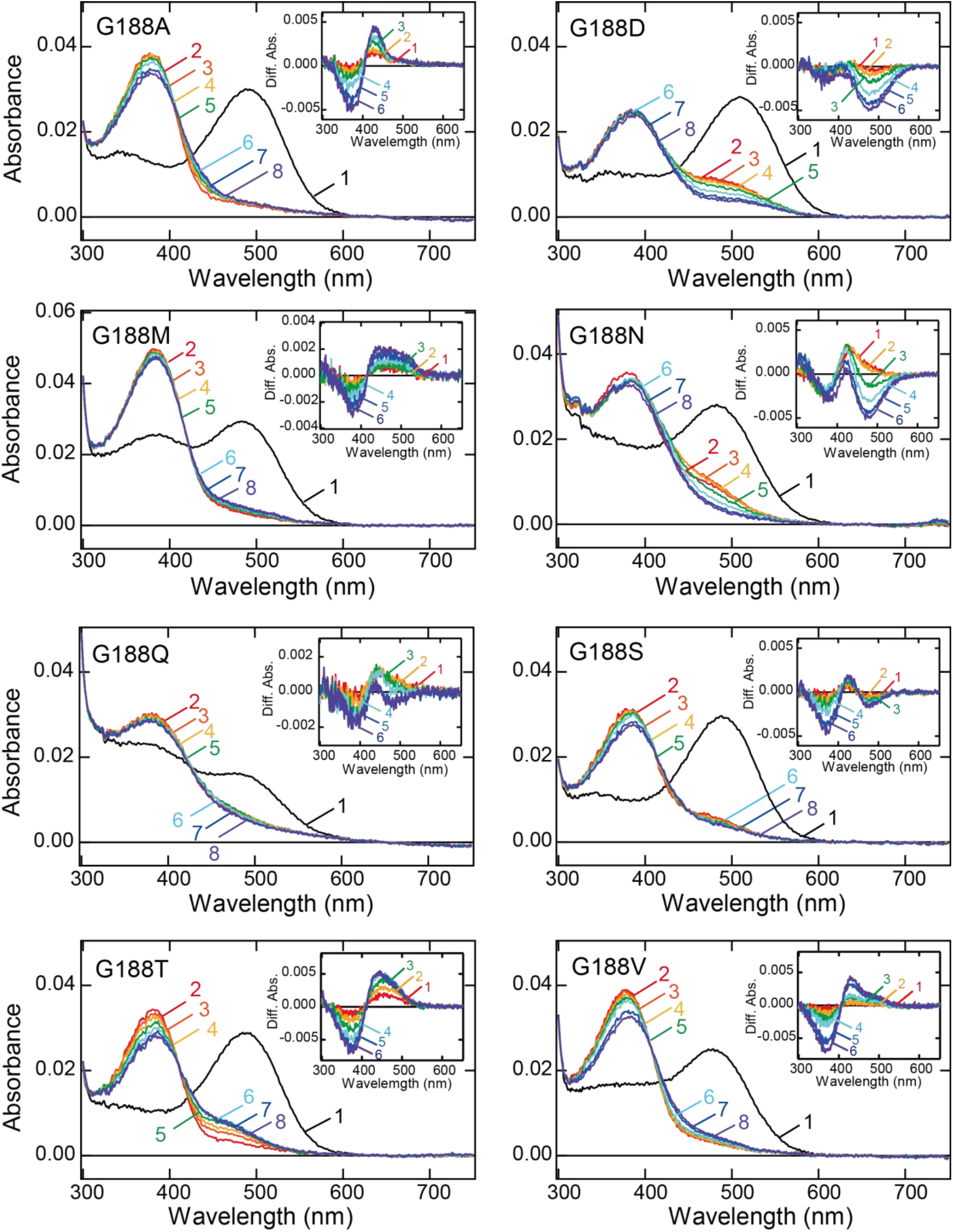
Thermal reaction of bovine rhodopsin Gly188 mutants after yellow light irradiation. Absorption spectra of G188A, G188D, G188M, G188N, G188Q, G188S, G188T and G188V mutants were recorded in the dark (curve 1) and 0, 5, 10, 20, 40, 80 and 120 min after yellow light (>500 nm) irradiation (curves 2-8, respectively) at 20 °C. (Inset) Difference spectra obtained by subtracting the spectrum just after irradiation (curve 2 in Fig. S3) from those measured after irradiation (curves 3-8 in Fig. S3) (curves 1-6, respectively).

**Figure S4.**
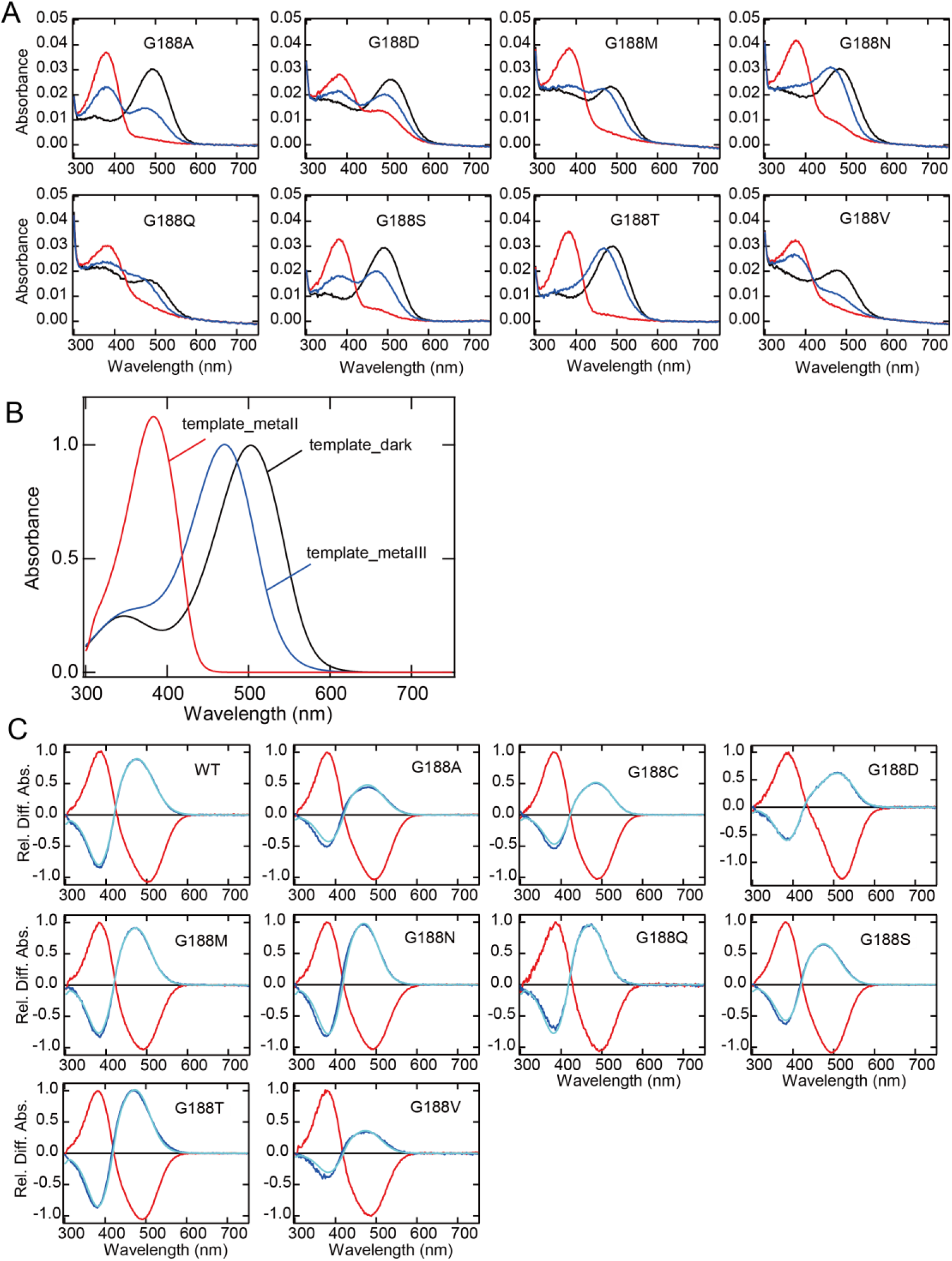
Absorption spectra of bovine rhodopsin Gly188 mutants. (A) Absorption spectra of G188A, G188D, G188M, G188N, G188Q, G188S, G188T and G188V mutants of bovine rhodopsin after reconstitution with 11-*cis* retinal. Spectra were recorded in the dark (black curve), after yellow light (>500 nm) irradiation (red curve) and after subsequent UV light (360 nm) irradiation (blue curve) at 0 °C. λmax of the mutants are shown in Table S1. (B) Template spectrum of the dark state, meta II and meta III of bovine rhodopsin wild-type modeled by the Lamb and Govardovskii method ^1, 2^. The template spectra of the dark state of mutants were constructed by the same method according to λmax of each mutant shown in Fig. S4A. The template spectrum of meta II was constructed from the absorption spectrum of native bovine rhodopsin irradiated at low pH. The template spectrum of meta III was constructed by the modification of the model spectrum of meta I which was constructed from the absorption spectrum of native bovine rhodopsin irradiated at high pH. The molar extinction coefficients of each state were taken from a previous report ^3^. (C) Spectral changes of the mutants induced by yellow light (red curve) and subsequent UV light (blue curve) irradiations at 0 °C. Difference spectra were calculated based on the spectra shown in Fig. S4A and were normalized to be ∼1.0 at the positive maximum of the red curve. The fitted curve (cyan curve) was obtained by fitting the blue curve using the template spectra of the dark state, meta II and meta III, and the component ratio of the dark state, meta II and meta III after UV light irradiation was calculated (Table S1).

**Figure S5.**
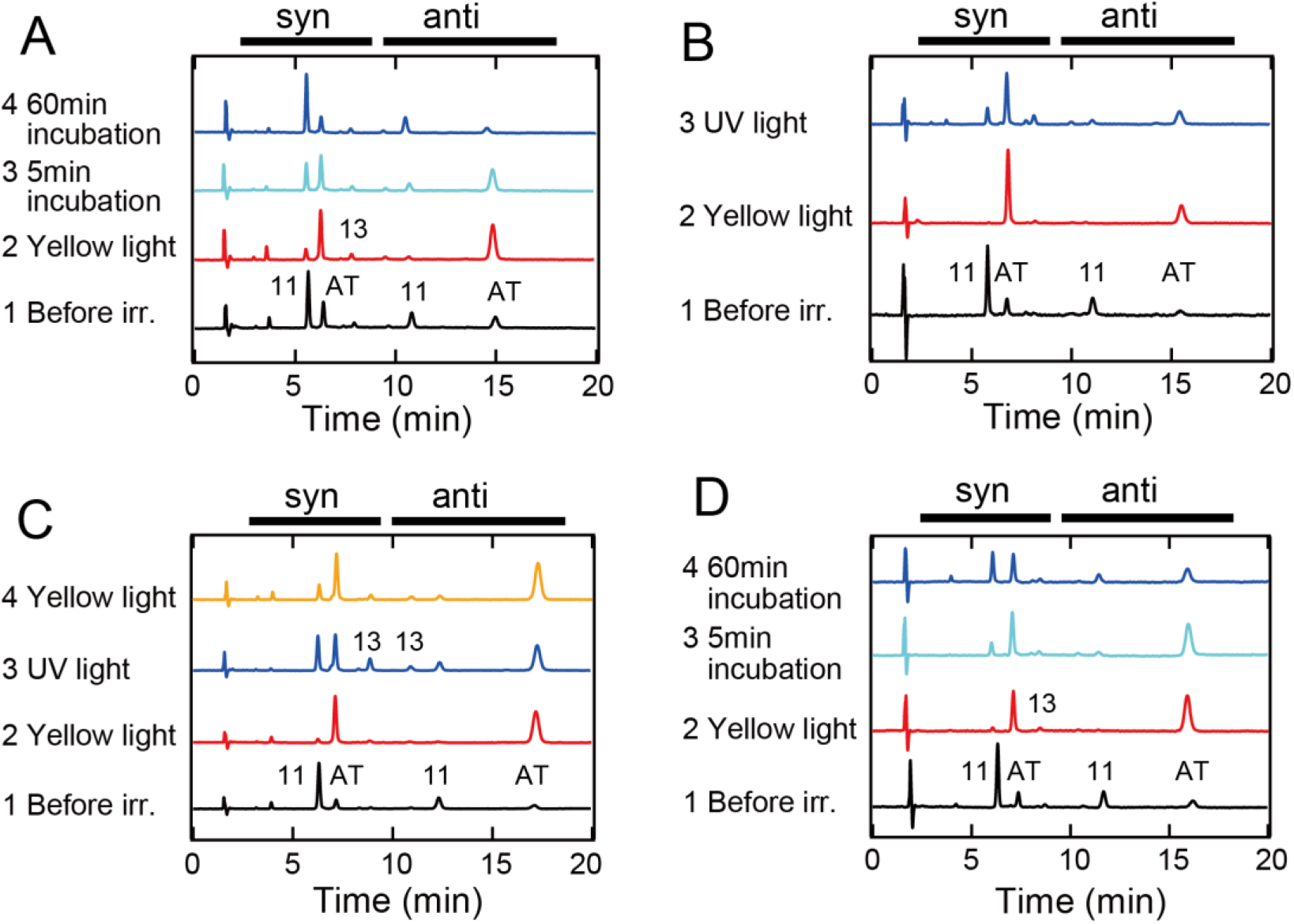
HPLC analysis of retinal configuration. (A) Bovine rhodopsin G188C/N2C/D282C mutant was purified after incubation with 11-*cis* retinal and was prepared for HPLC analysis of retinal configuration ^4^ 0, 5 and 60 min after yellow light (>500 nm) irradiation of the sample at 20 °C (see Figs. 1E and 1G). (B) Wild-type rhodopsin was purified after incubation with 11-*cis* retinal and was prepared for HPLC analysis after yellow light irradiation and subsequent UV light (360 nm) irradiation of the samples at 0 °C (see Figs. 2A and 2C). (C) G188C mutant was purified after incubation with 11-*cis* retinal and was prepared for HPLC analysis after yellow light irradiation, subsequent UV light irradiation and yellow light re-irradiation of the samples at 0°C (see Figs. 2B and 2D). (D) E122Q/G188C/N2C/D282C mutant was purified after incubation with 11-*cis* retinal and was prepared for HPLC analysis 0, 5 and 60 min after yellow light irradiation of the sample at 0 °C (see Figs. 3A and 3B).

**Figure S6.**
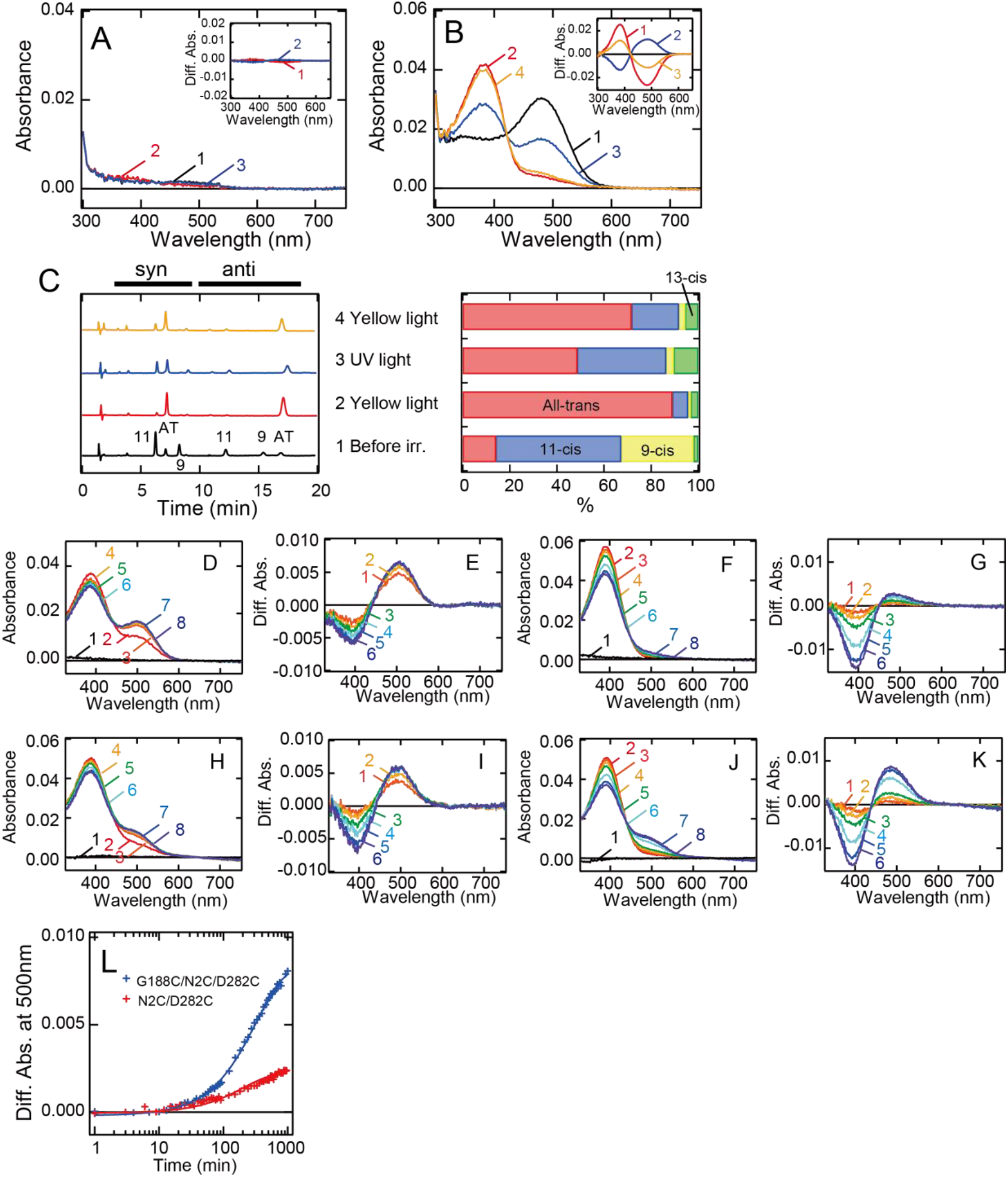
Formation of the photo-pigments of bovine rhodopsin G188C mutant after incubation with all-*trans* retinal. (A, B) Absorption spectra of wild-type (A) or G188C mutant (B) purified after the addition of all-*trans* retinal to the suspension of rhodopsin-expressing cell membranes at 0 °C. Spectra were measured in the dark (curve 1), after yellow light (>500 nm) irradiation (curve 2), after subsequent UV light (360 nm) irradiation (curve 3) and after yellow light re-irradiation (curve 4). (Inset) Spectral change caused by yellow light irradiation (curve 1), subsequent UV light irradiation (curve 2) and yellow light re-irradiation (curve 3). (C) Isomeric compositions of retinal of G188C mutant purified after the addition of all-*trans* retinal. (*Left*) Retinal configurations were analyzed by HPLC analysis after extraction of the chromophore as retinal oximes from the samples before light irradiation, after yellow light irradiation, subsequent UV light irradiation and yellow light re-irradiation. (*Right*) Changes of retinal isomer compositions of G188C mutant. (D-G) Regeneration of the photo-pigments by the addition of 11-*cis* (D, E) or all-*trans* (F, G) retinal to purified apo-protein of N2C/D282C. (D) Spectra were measured before (curve 1) and 0, 3, 6, 15, 30, 60 and 120 min after the addition of 1.1 μM 11-*cis* retinal (curves 2-8). (E) Difference spectra were calculated by subtracting the spectrum just after the addition of 11-*cis* retinal (curve 2 in Fig. S6D) from those measured 3, 6, 15, 30, 60 and 120 min after the addition of 11-*cis* retinal (curves 3-8 in Fig. S6D) (curves 1-6, respectively). (F) Spectra were measured before (curve 1) and 0, 0.5, 1, 2, 6, 12 and 16 hours after the addition of 1.1 μM all-*trans* retinal (curves 2-8). (G) Difference spectra were calculated by subtracting the spectrum just after the addition of all-*trans* retinal (curve 2 in Fig. S6F) from those measured 0.5, 1, 2, 6, 12 and 16 hours after the addition of all-*trans* retinal (curves 3-8 in Fig. S6F) (curves 1-6, respectively). (H-K) Regeneration of the photo-pigments by the addition of 11-*cis* (H, I) or all-*trans* (J, K) retinal to purified apo-protein of G188C/N2C/D282C. (H) Spectra were measured before (curve 1) and 0, 3, 6, 15, 30, 60 and 120 min after the addition of 1.1 μM 11-*cis* retinal (curves 2-8). (I) Difference spectra were calculated by subtracting the spectrum just after the addition of 11-*cis* retinal (curve 2 in Fig. S6H) from those measured 3, 6, 15, 30, 60 and 120 min after the addition of 11-*cis* retinal (curves 3-8 in Fig. S6H) (curves 1-6, respectively). (J) Spectra were measured before (curve 1) and 0, 0.5, 1, 2, 6, 12 and 16 hours after the addition of 1.1 μM all-*trans* retinal (curves 2-8). (K) Difference spectra were calculated by subtracting the spectrum just after the addition of all-*trans* retinal (curve 2 in Fig. S6J) from those measured 0.5, 1, 2, 6, 12 and 16 hours after the addition of all-*trans* retinal (curves 3-8 in Fig. S6J) (curves 1-6, respectively). (L) Regeneration processes of the photo-pigments of N2C/D282C (red curve) and G188C/N2C/D282C (blue curve) by the addition of all-*trans* retinal as shown in Figs. S6F and S6J were monitored by the change of absorbance at 500nm.

**Figure S7.**
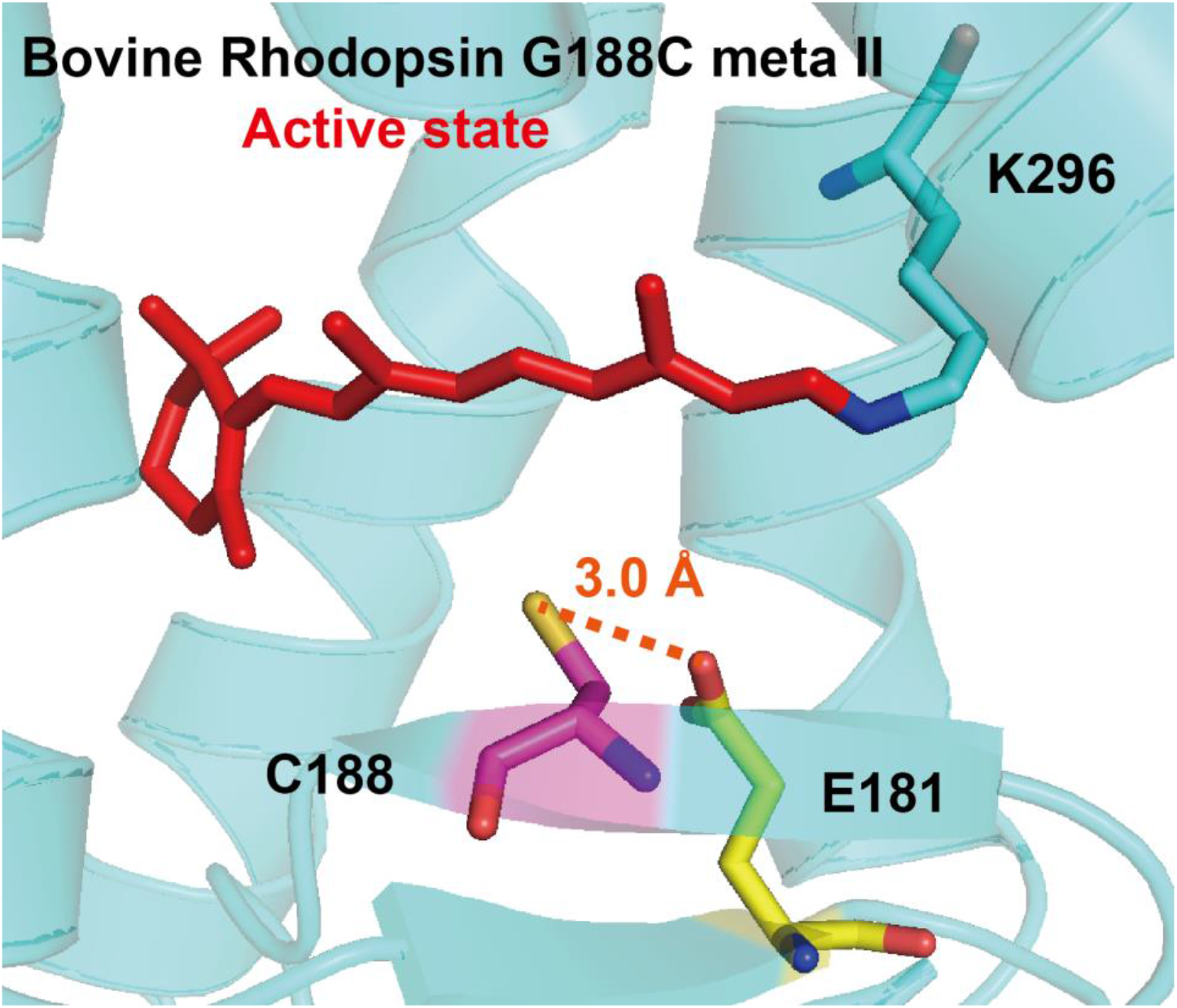
Predicted structural model around position 188 of G188C mutant meta II. The structural model of G188C mutant meta II was constructed based on the 3D structure of wild-type meta II ^5^ using PyMOL. Cys188 can be located in the vicinity of Glu181 which serves as a counterion to stabilize the protonation of the Schiff base in meta I, a precursor of meta II ^6, 7^.

**Figure S8.**
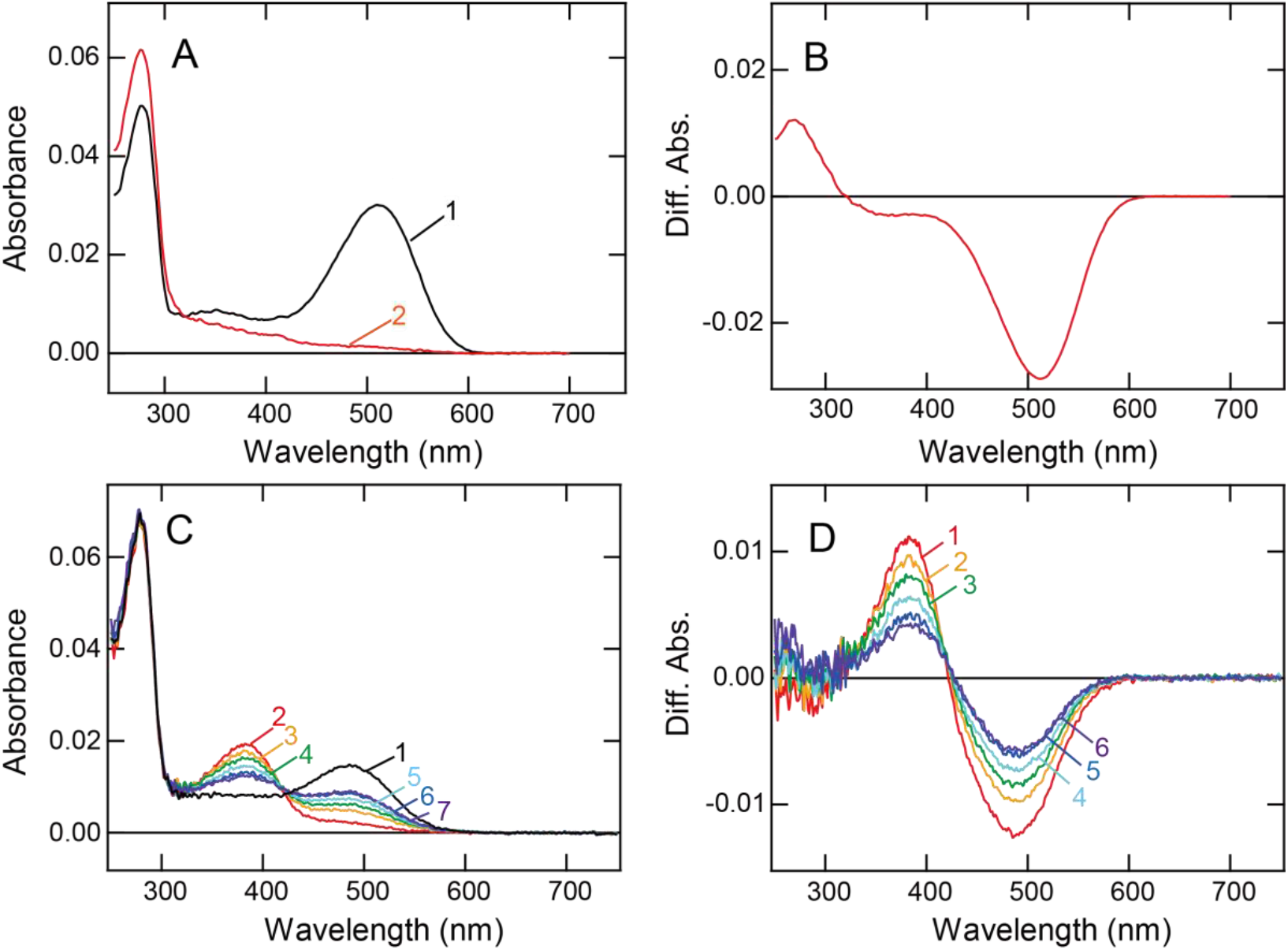
Absorption spectra of Opn5L1 and bovine rhodopsin G188C mutant. (A, B) Spectral change of chicken Opn5L1. (A) Spectra were recorded in the dark (curve 1) and after yellow light (>500 nm) irradiation (curve 2) as referred to our previous paper ^8^. (B) Difference spectrum was obtained by subtracting the spectrum before irradiation (curve 1 in Fig. S8A) from the spectrum measured after irradiation (curves 2 in Fig. S8A). A light-dependent substantial increase of the absorbance was observed at around 270 nm. (C, D) Spectral changes of bovine rhodopsin G188C mutant. (C) Spectra were recorded at 20 °C in the dark (curve 1) and 0, 5, 10, 20, 40 and 80 min after yellow light (>500 nm) irradiation (curves 2-7, respectively). (D) Difference spectra were obtained by subtracting the spectrum before irradiation (curve 1 in Fig. S8C) from the spectra measured after irradiation (curves 2-7 in Fig. S8C) (curves 1-6, respectively). Absorbance changes were observed in the visible and near-UV regions (350∼700 nm), not at around 270 nm.

**Table S1.** Comparison of λmax in the dark state and spectral components after UV light irradiation.

| | $\lambda_{\text{max}}^{\text{a}}$ | dark state<sup>b</sup> (%) | meta II<sup>b</sup> (%) | meta III<sup>b</sup> (%) |
| --- | --- | --- | --- | --- |
| wild-type | 500 | 21.6<sup>c</sup> | 13.7 | 64.7 |
| G188C | 487 | 41.2<sup>c</sup> | 48.5 | 10.3 |
| G188A | 494 | 18.1 | 51.7 | 30.2 |
| G188D | 509 | 30.9 | 67.7 | 1.4 |
| G188M | 491 | 9.2 | 10.0 | 80.8 |
| G188N | 492 | 0 | 0 | 100 |
| G188Q | 493 | 2.4 | 5.5 | 92.1 |
| G188S | 495 | 14.6 | 37.7 | 47.7 |
| G188T | 488 | 11.0 | 2.9 | 86.1 |
| G188V | 486 | 7.3 | 63.4 | 29.3 |

<sup>a</sup> $\lambda_{\text{max}}$ was estimated from the absorption spectra shown in Figs. 2 and S3.
<sup>b</sup>Component ratios of the dark state, meta II and meta III were calculated based on the spectral changes induced by UV light irradiation shown in Figs. 2 and S3.
<sup>c</sup>Ratios of the dark state in wild-type and G188C are comparable to those of 11-*cis* retinal obtained by the retinal configuration analysis (24% in wild-type and 40 % in G188C).

